# A multi-parameter optimization in middle-down analysis of monoclonal antibodies by LC-MS/MS

**DOI:** 10.1101/2022.12.08.518878

**Authors:** Jonathan Dhenin, Mathieu Dupré, Karen Druart, Alain Krick, Christine Mauriac, Julia Chamot-Rooke

**Affiliations:** Mass Spectrometry for Biology Unit, Université Paris Cité, Institut Pasteur, CNRS UAR2024, Paris, France; DMPK, Sanofi, Chilly-Mazarin, France; Université Paris Cité, Sorbonne Paris Cité, Paris, France

## Abstract

In antibody-based drug research, regulatory agencies request a complete characterization of antibody proteoforms covering both the amino acid sequence and all post-translational modifications. The usual mass spectrometry-based approach to achieve this goal is bottom-up proteomics, which relies on the digestion of antibodies, but does not allow the diversity of proteoforms to be assessed. Middle-down and top-down approaches have recently emerged as attractive alternatives but are not yet mastered and thus used in routine by many analytical chemistry laboratories. The work described here aims at providing guidelines to achieve the best sequence coverage for the fragmentation of intact light and heavy chains generated from a simple reduction of intact antibodies using Orbitrap mass spectrometry. Three parameters were found crucial to this aim: the use of an electron-based activation technique, the multiplex selection of precursor ions of different charge states and the combination of replicates.

## 1. INTRODUCTION

In the last twenty years, antibodies (Abs) have become the new backbone of the pharmaceutical industry. The use of monoclonal Abs (mAbs) is now approved for over 67 targets, mostly involved in oncology but not only^1,2^. Because mAbs are large molecules, their characterization requires the development of specific mass spectrometry approaches. Bottom-up proteomics (BUP), which is based on the digestion of Abs often with multiple enzymes, is the primary line of analysis, allowing both sequence information and post-translational modification localization (PTMs) to be achieved in many cases^3–6^. However, this approach does not bring the required information on disulfide bond arrangements, or simply on the different proteoforms present in a sample. These proteoforms can arise from N-terminal pyroglutamic acid formation, C-terminal clipping, glycosylation, glycation, methionine oxidation, deamidation, cysteinylation, etc^7^. To fulfill this goal, alternative and complementary approaches have emerged, either on the intact mAbs (top-down proteomics or TDP) or on large parts of the antibodies obtained either by S-S reduction or by specific enzymatic digestion^8–11^ (middle-down proteomics or MDP). A recent interlaboratory study on the characterization of intact antibodies by TDP and MDP approaches including 20 partner laboratories and a wide diversity of instrumentation and methods highlighted the fact that there is no “one size fits all”^12^. One of the conclusions of this study is that compared to the techniques and instruments used for BUP, TDP and MDP show a clear potential for high performance protein analysis but still require developments. One of the main advantages of TDP/MDP approaches is that the information achieved from MS/MS spectra is complementary to the BUP results and can also be obtained more rapidly. A very important piece of information is the molecular weight of intact mAbs. For extensive MS/MS characterization, the size and structure of these molecules (with both the intra- and inter-chain S-S bonds) lead to incomplete sequence coverage that could be problematic to cover properly all Complementary Determining Regions (CDRs)^13^. Combining several activation techniques such as collision-based (collision-induced dissociation, CID; higher-energy C-trap dissociation, HCD), electron-based (electron-transfer dissociation, ETD; electron-capture dissociation, ECD), combination of thereof (electron-transfer supplemented with collision or higher-energy collision dissociation, ETciD or EThcD) or photon-based (ultraviolet photodissociation, UVPD) improves this issue but never leads to a 100% coverage^13–17^. One strategy, developed by several groups, is to focus on the analysis of antibody subunits which can be either the light chain (Lc) and heavy chain (Hc) after disulfide reduction^18,19^, Fab and Fc after specific enzymatic digestion^13,20–22^ or Lc, Fd and Fc/2 when combining both approaches^13,23–27^. However, even with the smallest 25 kDa subunits a single MS/MS acquisition does not yield the expected sequence coverage and a combination of experiments are required. To date, no generic approach has been developed for this family of molecules, although such approach would be of high interest for biopharma companies. The objective of this paper is to provide a set of optimized instrumental parameters for the sequencing of Lc and Hc by LC-MS/MS on an Orbitrap Tribrid mass spectrometer. Indeed, in contrast to BUP where the number of factors to optimize are very limited, in TDP/MDP the quality of results strongly depends on a few parameters such as the precursor charge state^23,28–33^, the activation technique and level (energy or time)^6,13,18,19,23–27,29^ and number of replicates. Compared to infusion experiments, the chromatographic separation brings additional constraints that will be discussed. Our results highlight the importance of electron-based fragmentation for improved sequence coverage. Initially restricted to a certain class of mass spectrometers (ECD in FT-ICR MS, ETD in ion traps), these methods are now more widely available on diverse platforms.

## 2. MATERIAL AND METHODS

### 2.1. Antibody sample preparation

Two commercially available mAbs (immunoglobulin or IgG, type 1) were used for the fragmentation experiments: Sigma mAb standard (SiLuLite, IgG1 lambda, CHO, Sigma) and NIST mAb standard (HzIgG1 kappa, NS0, NIST). These mAbs were chosen based on the strong sequence differences between their light chains (only 46% identity based on sequence alignment). Each mAb was diluted to a final concentration of 0.32 μg/μL with guanidine hydrochloride (Sigma, 5 M final concentration) and reduced into Lc and Hc with DTT (Sigma, 100 mM final concentration) during 45 min at 45°C under 800 rpm. Samples were acidified with TFA (Sigma, 1% final concentration) before LC-MS/MS analysis.

### 2.2. Liquid chromatography – mass spectrometry

LC-MS/MS analysis was performed using a Vanquish Horizon UHPLC system (Thermo Scientific, San Jose, CA) coupled to an Orbitrap Eclipse™ Tribrid mass spectrometer (Thermo Scientific, San Jose, CA) fitted with a H-ESI source. 1 μg of Lc and Hc were separated on a MAbPac™ RP column (2.1 mm i.d. x 50 mm, 4 μm particle size, 1500 Å pore size, Thermo Scientific, San Jose, CA) heated at 80°C with a gradient of 0.1% formic acid in acetonitrile (ACN) at 250 μL/min, from 25 to 41% in 2.8 min. For all experiments spray voltage was set to 3.8 kV, sheath gas settings was 35, auxiliary gas settings was 4, sweep gas settings was 1, vaporizer temperature was 150°C, ion transfer tube temperature was 350°C, RF value was 30% and source fragmentation energy was 10 V. A first LC-MS experiment was acquired at 7,500 resolving power (at m/z 400) with a scan range set to m/z 350-2,500, 5 microscans (μscans) per MS scan, an automatic gain control (AGC) target value of 5×10^5^, a maximum injection time of 50 ms. Fragmentation data were recorded using targeted LC-MS/MS experiments between 2.5 and 3.2 min for the NIST mAb Lc and 3.2 and 4.2 min for the NIST mAb Hc; between 2.6 and 3.25 min for the Sigma mAb Lc and 3.2 and 4.0 min for the Sigma mAb Hc. Four precursor charge states were chosen for each subunit across their respective charge state distribution, isolated by the quadrupole and subjected to individual or multiplexed fragmentation (Table S1). MS/MS scans were acquired at 120,000 resolving power (at m/z 400) with an isolation width of 1.6 m/z, 5 μscans, an AGC target value of 5×10^5^ and maximum injection time of 246 ms. HCD with normalized collision energies (NCE) of 15, 20 and 25%, EThcD with 2, 5 and 10 ms of reaction time and a supplemental HCD activation with NCE of 5, 10 or 15%, CID with collision energies (CE) of respectively 25, 30, 35% and 30, 35, 40% were used for the fragmentation of the Lc and Hc. For EThcD experiments, the anionic fluoranthene reagent AGC target was set to 7×10^5^ with a maximum injection time of 200 ms. All experiments were conducted using the Intact Protein mode with a pressure set to 1 mTorr in the ion-routing multipole (IRM).

### 2.3. Data analysis

MS spectra were deconvoluted with Genedata Expressionist^®^ software using a time-resolved deconvolution and the Maximum Entropy (MaxEnt) algorithm. MS/MS spectra were averaged across the appropriate subunit elution windows and then deconvoluted using the embedded Xtract algorithm in FreeStyle™ (v. 1.6.75.20) with a signal-to-noise ratio threshold of 3, a fit factor of 80%, a remainder threshold of 25% and maximum charge set to the precursor charge state. Lists of decharged and deisotoped monoisotopic masses were imported into ProSight Lite (v. 1.4)^34^ and used for fragment assignments with a 5 ppm mass tolerance. Only b- and y-ions were considered for CID and HCD fragmentations while b-, c-, y- and z-ions were searched for EThcD. TDFragMapper^35^ was used for further visualization and comparison of fragmentation results. Lists of assigned fragments were exported from ProSight Lite and used with deconvoluted data and the protein sequence as input for TDFragMapper. Venn diagrams and pairwise comparisons were generated using the Intervene Shiny app^36^. Combinations of fragmentation results from diverse MS/MS experiments were processed using an in-house developed R script.

## 3. RESULT AND DISCUSSION

### 3.1. Precursor charge state

Ionization of the reduced Lc and Hc from both NIST and Sigma mAbs in denaturing conditions yields two charge state envelopes from 10+ to 30+ for the Lc and from 30+ to 65+ for the Hc (Figure 1 for NISTmAb and figure S1 for Sigma mAb).

**Figure 1.**
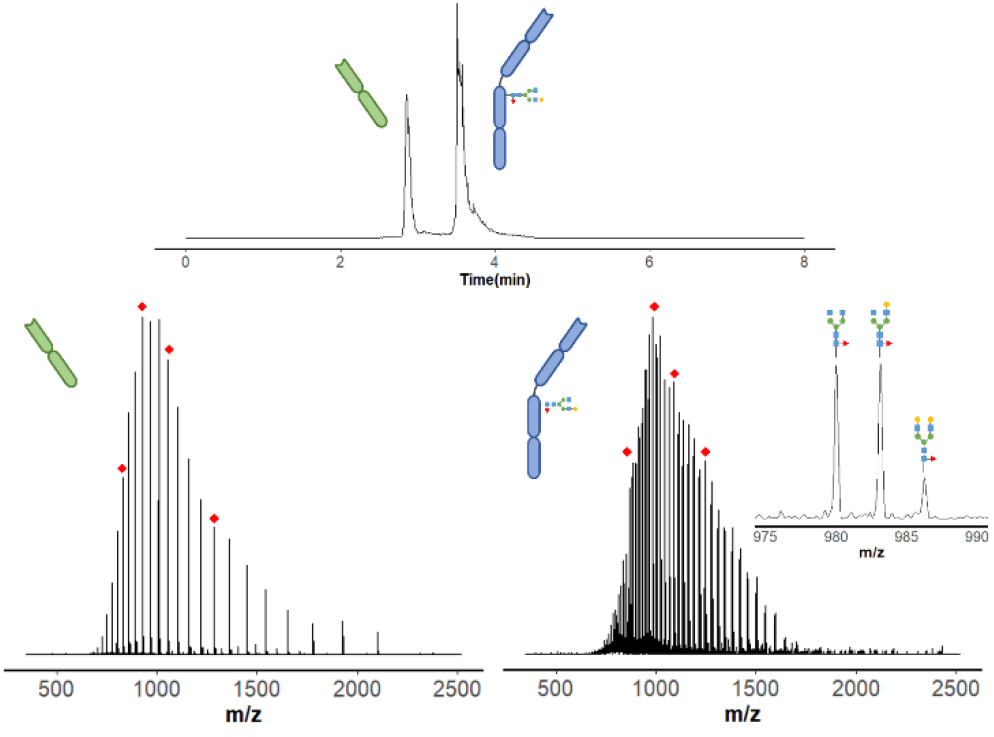
Chromatogram of NISTmAb Lc and Hc (top), MS spectra of Lc (bottom left) and Hc (bottom right). Precursor charge states selected for MS/MS analysis are depicted as red diamonds in MS spectra.

From this envelope, the most abundant charge state or several consecutive abundant charge states with a large isolation window are often selected for MS/MS fragmentation. Co-elution of multiple proteoforms in LC-MS can however hinder the use of a wide isolation window to avoid their simultaneous fragmentation and the production of chimeric spectra. In recent instruments, individual charge states can be either individually selected with a very narrow window, or in a multiplexed manner to improve signal/noise ratio of the fragment ions. In top-down mass spectrometry, a clear link between the precursor charge state and the quality of the MS/MS spectra has already been described ^23,28–33^ and thus this parameter should be carefully optimized. To evaluate the role of this parameter when fragmentating Lc and Hc, 4 different charge states centered on the most abundant one in the envelope were selected either individually or simultaneously (multiplex). The lowest charge states (below 15+ for the Lc and below 40+ for the Hc) and highest ones were excluded from the study to avoid MS/MS spectra with very low intensity and competitive background noise. For the Hc, the three major glycoforms were co-selected for each charge state (Figure 1, right inset). As expected, very different raw and deconvoluted MS/MS spectra were obtained for the HCD fragmentation of both NISTmAb Lc and Hc (Figure 2 and figure S2). Similar results were obtained with other fragmentation techniques. As depicted on Figure 2, shifting from 17+ to 27+ (or multiplex) leads to very different fragmentation patterns, with a different distribution of fragment ions along both the mass and intensity axes. HCD activation of highly charged precursor ions generates more smaller fragments than the activation of lower charged precursors with the same activation energy. Less golden complementary pairs are also obtained when fragmenting highly charged precursors. The smallest fragments (low mass region) can correspond to the cleavage of both the N- and C-terminal extremities and are therefore of interest to increase the sequence coverage (Figure S3). Although many of the observed fragments are common to all precursor charge states, unique ones can be retrieved for each individual precursor (both for the Lc and Hc subunits). Most of these unique fragments can be recovered in the multiplex experiments for the Lc (between 79 and 93%), but much less for the Hc (between 30 and 90%) (Table 1 and Table S2).

**Table 1.** Comparison of residue cleavages between multiplex and the sum of individual precursors for the NISTmAb Lc and percentages of cleavages from individual fragmentations contained in multiplex experiments

| Fragmentation experiment | Cumulated residue cleavages (%) |  | Cleavages from individual precursors observed in multiplex (%) |
| --- | --- | --- | --- |
|  | Multiplex | Individual precursors |  |
| HCD 15% | 30 | 33 | 86 |
| HCD 20% | 34 | 35 | 91 |
| HCD 25% | 34 | 37 | 87 |
| ET2hcD5 | 65 | 77 | 80 |
| ET2hcD15 | 63 | 77 | 79 |
| ET5hcD5 | 74 | 79 | 90 |
| ET5hcD15 | 75 | 74 | 92 |
| ET10hcD10 | 73 | 66 | 93 |
| ET10hcD15 | 68 | 58 | 92 |

**Figure 2.**
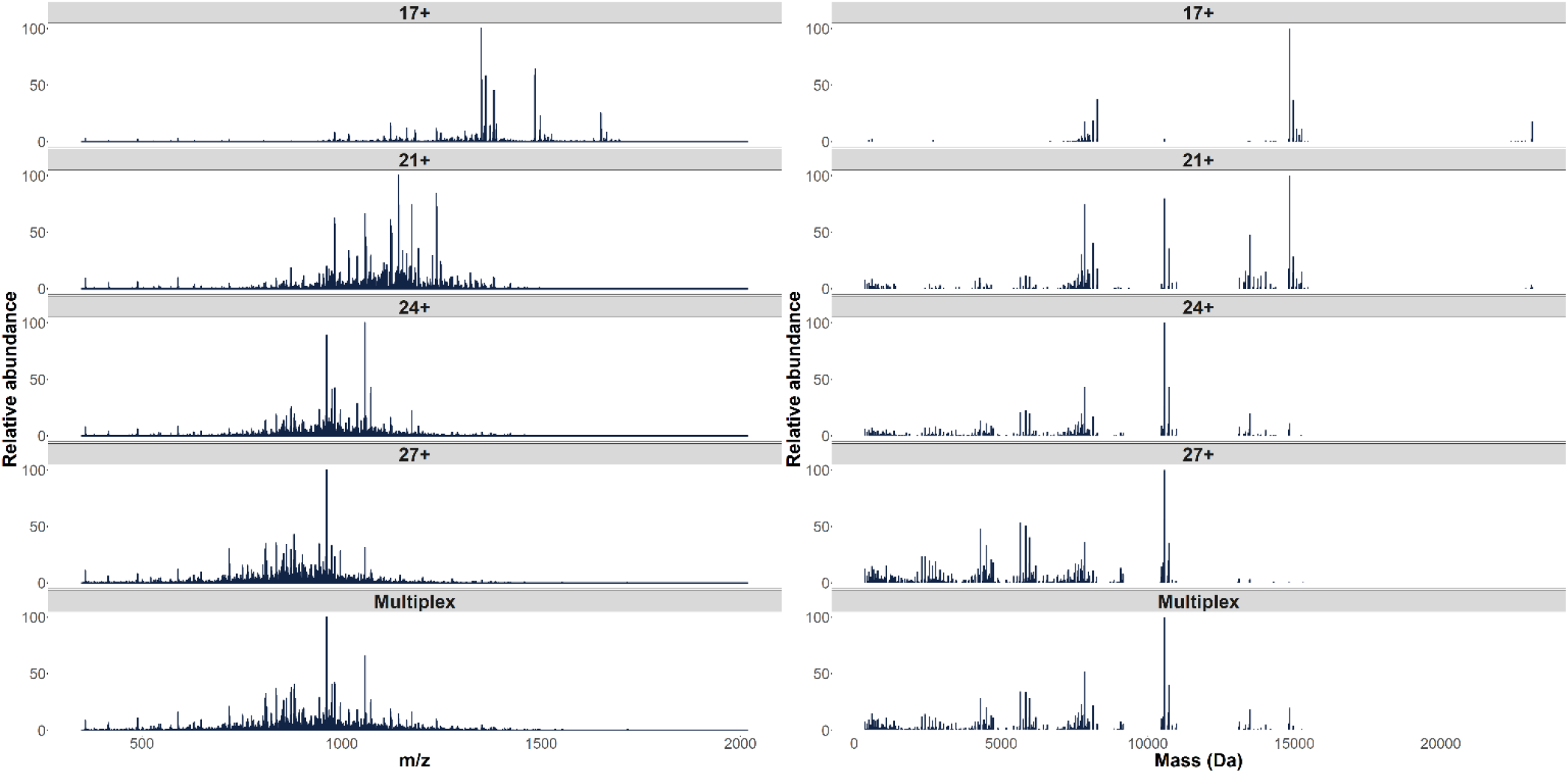
HCD MS/MS spectra with NCE25% for each selected charge state of NISTmAb Lc before (left) and after deconvolution (right). From top to bottom: 17+, 21+, 24+, 27+ and multiplex

Multiplexing several precursor charge states for the Lc analysis seems therefore to be a valuable strategy to combine most of the unique fragments of individual precursors into a single fragmentation experiment. This result obtained for a single replicate is also true for a triplicate: 3 multiplex experiments are almost equivalent to 12 individual ones (4 precursors x 3 replicates) for HCD but can also give better results for EThcD (Table 1). For the Hc, the situation is slightly different and the sum of the 12 individual experiments always outperforms the multiplex ones for all tested HCD energies as well as for short EThcD experiments (Table S2). Note however that we compare here the results of 12 experiments (individual precursors) to those obtained only for 3 experiments (multiplex). Our results highlight that targeting different precursor charge states can provide complementary sequence information as different fragmentation pathways are activated. Moreover, using a multiplex selection is highly attractive as it reduces the number of experiments required to achieve the best sequence coverage.

### 3.2. Fragmentation method

CID and HCD were compared to EThcD (Figure 3). We decided not to include ETD because its efficiency as a standalone technique has already proved limited for the fragmentation of intact proteins, especially in an LC time scale. ETD is a low-efficiency activation technique, which has to be combined with supplementary activation to yield satisfactory results^30,37,38^. The data obtained on the NISTmAb Lc indicate that both CID and HCD give similar results with a sequence coverage of 35% maximum. EThcD gives the best results in all cases (up to 66% in a single experiment), independently of the precursor charge state. Moreover, although multiplexing does not really bring a substantial improvement of CID/HCD, this is not the case for EThcD especially with the largest activation times (5-10 ms). A similar conclusion can be drawn for the analysis of the NISTmAb Hc (Figure S4), with EThcD leading to the best results despite a lower sequence coverage (maximum 19% for the multiplex in a single experiment) than what was achieved for the Lc. Similar results were obtained with the Sigma mAb standard (Figures S5 and S6). We then compared the results obtained using each fragmentation method for the same precursor charge state (Figures 4 for NISTmAb Lc and S7 for NISTmAb Hc). The Venn diagrams confirm that EThcD brings additional and unique cleavages compared to CID and HCD (although slower), with a charge state-dependency. For the fragmentation of the NISTmAb Lc, the percentage of unique cleavages provided by EThcD ranges from 37% to 46% for individual precursor ions, and up to 67% for the multiplex. This is particularly the case for the higher charge states and less pronounced for the lowest ones for which CID leads to a significant number of unique cleavages.

**Figure 3.**
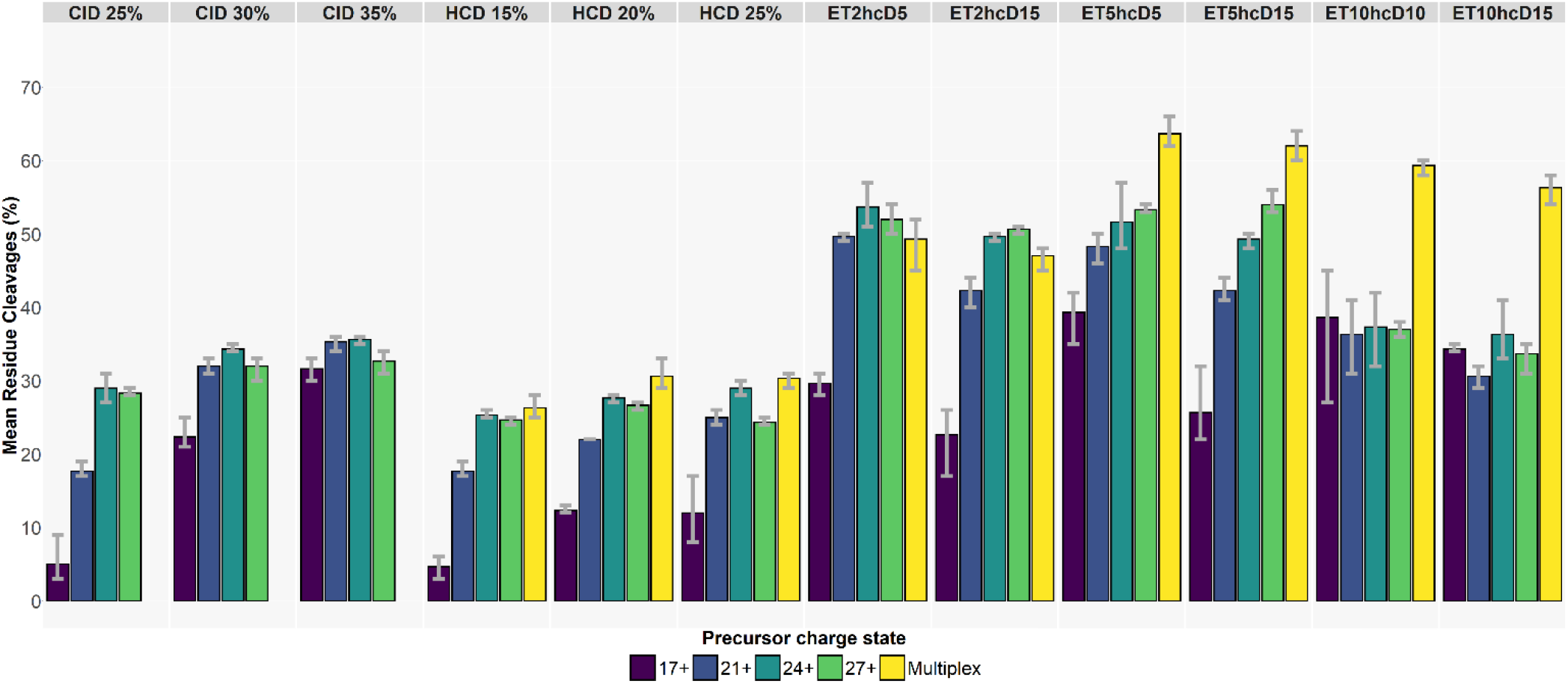
Mean residue cleavages (triplicates) for each selected precursor charge states of the NISTmAb Lc across the different fragmentation experiments

**Figure 4.**
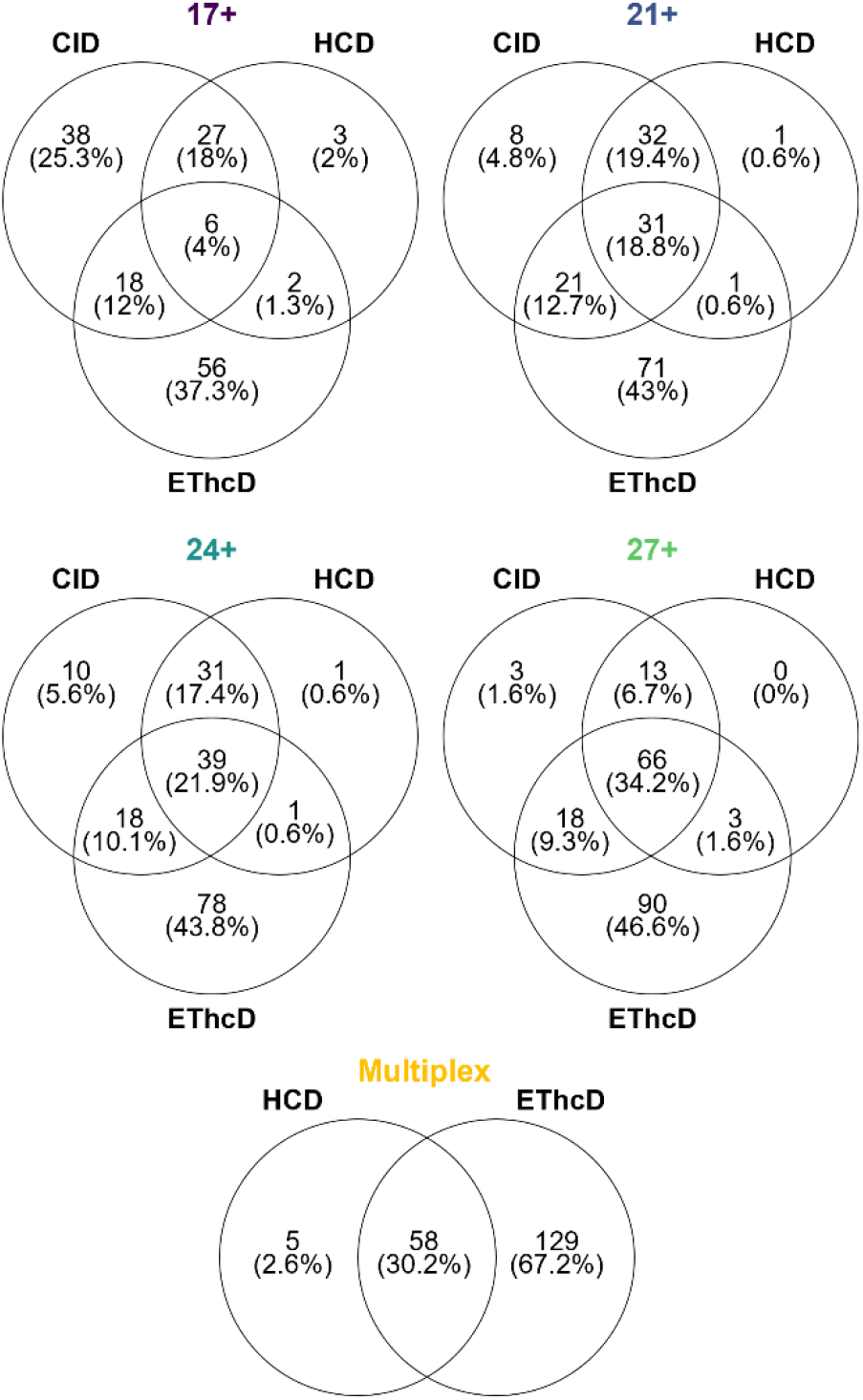
Venn diagrams of cleavages generated in CID, HCD and EThcD for every targeted precursor charge state and multiplex for the NISTmAb Lc

These unique cleavages represent 25% of all cleavages for the 17+ of the Lc but only 10% for the 40+ of the Hc, which is the lowest charge state fragmented. In contrast to CID, HCD on the Hc also brings unique cleavages (11-18%) that can be explained by the different energetic regime of both techniques. Concerning the general overlap of all three fragmentation techniques, the maximum achieved is 34% for the Lc and 8% for the Hc, which strongly supports the combination of fragmentation techniques for an improved sequence coverage. In conclusion, EThcD should always been favored since it leads to numerous unique fragments that cannot be obtained in HCD or CID.

### 3.3. Technical replicates

Lc and Hc fragmentations take place on a chromatographic time scale and therefore the number of scans that can be accumulated is limited by the chromatographic peak width, which is 6 s in our conditions. Within this timeframe and with a resolution set to 120k, a maximum of 30 microscans can be acquired per LC run. Fragmentation of intact proteins with molecular masses above 25 kDa requires the accumulation of the largest number of scans to allow low intensity fragments to be extracted from the background noise. This scan accumulation remains one of the biggest challenges in LC-MS/MS experiments (compared to fragmentations acquired using infusion). One possible strategy to cope with this issue is to perform several replicates through repeated injections. Three replicates were therefore acquired for each fragmentation experiment in our study to assess the fragmentation repeatability and the expected added value of replicates. Cumulated percentages of residue cleavages were computed and summarized for the NISTmAb Lc and Hc as shown in Figure 5 and Figure S8, respectively. Our results indicate that increasing the number of replicates leads to a significant improvement for EThcD (up to 23% for the Lc and 11% for the Hc with ET5hcD15) but much less for CID and HCD (maximum 9%/5% for Lc/Hc). The maximum sequence coverage obtained for 3 EThcD replicates is 75% for the Lc and 27% for the Hc, to be compared to 66% and 16%, respectively, for a single replicate. The improvement is therefore, as expected, more pronounced for the Hc. Similar observations could be made for the second mAb studied (Figures S9 and S10). The difference observed between EThcD and CID/HCD can probably be explained by the fact that EThcD opens more fragmentation channels than CID or HCD (with the formation of both c/z and b/y ions and not only b/y ones) and thus the fragment ions formed are of lower intensity. Cumulating replicates allows the diversity of EThcD cleavages to be better explored. It is also worth noting that the repeatability of the fragmentation is slightly influenced by the precursor charge state (and inherently by its relative intensity in the whole charge envelope). The conclusion here is that cumulating at least 2 technical replicates, ideally 3, improves the global sequence coverage of both Lc and Hc.

**Figure 5.**
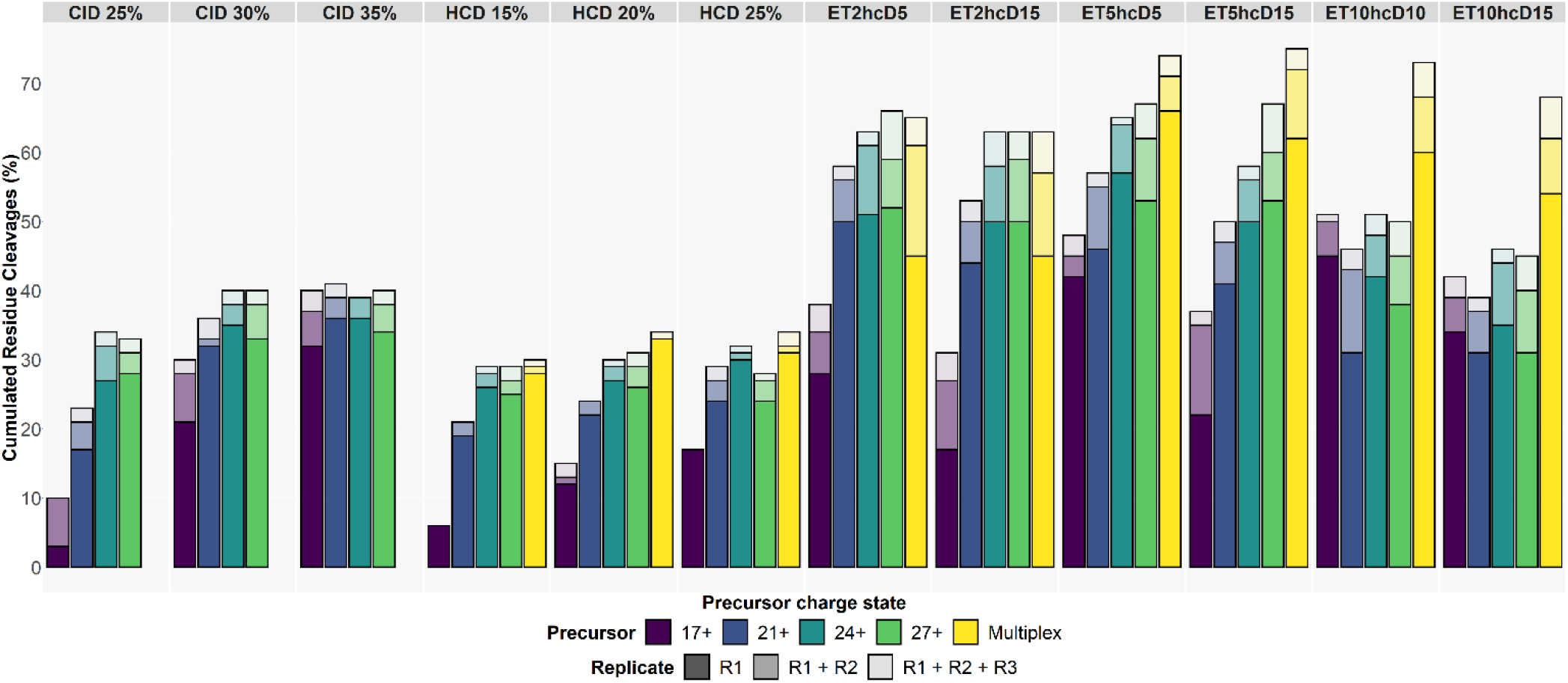
Summed residue cleavages over 3 replicates for each selected precursor charge states of the NISTmAb Lc across the different fragmentation experiments

### 3.4. Optimized combination of parameters

The objective here was to evaluate which combination of parameters, associated with the lowest number of experiments, can achieve the best sequence coverage for both Lc and Hc. In total we performed 57 different experiments (in triplicate) for each subunit, varying the precursor charge state, the fragmentation type and the activation level. The pairwise combination of all conditions would lead to 2^57^ possibilities (Figure S11), which is impossible to compute. We therefore decided to take the 3 best sets of parameters of each charge state and proceed with the comparison. The number of possibilities is now reduced to 2^15^ (32,768) which can be easily handled. The results are summarized in Figure 6 for NISTmAb (Figure S12 for Sigma mAb).

**Figure 6.**
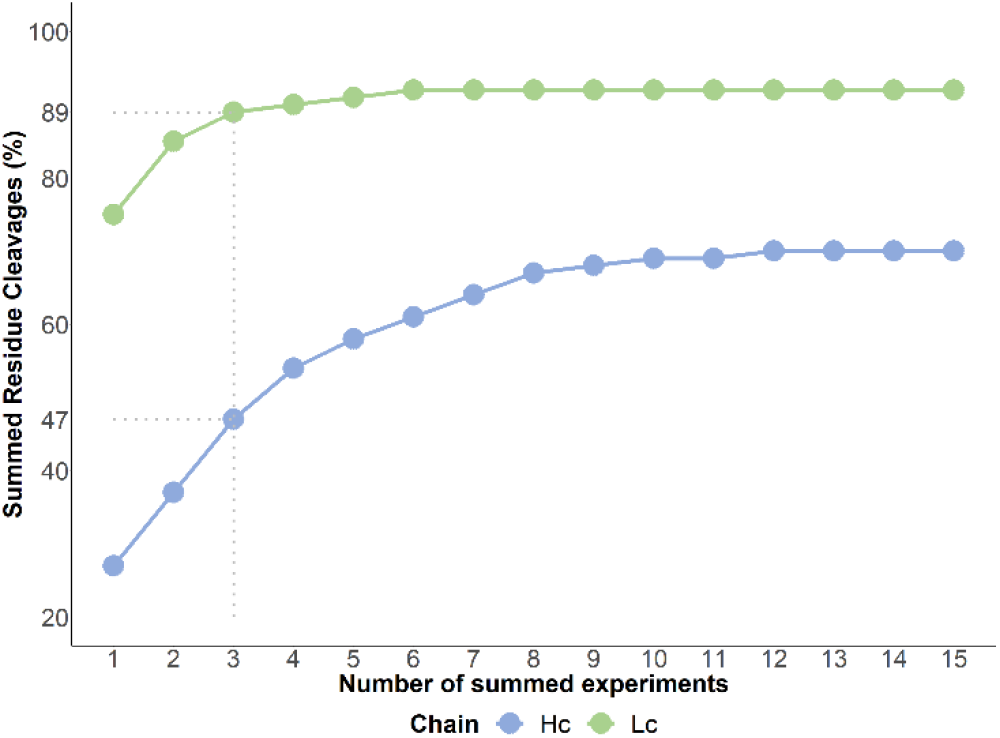
Maximum residue cleavage (%) obtained when combining diverse MS/MS experiments for the NISTmAb Lc and Hc

In line with the results described above, the combination of multiplex experiments always outperforms the others, whatever the types and level of fragmentation used, for both Lc and Hc. The best combination of 3 LC-MS/MS runs is composed by two EThcD experiments and one HCD. This shows that HCD still brings unique fragments that cannot be obtained by EThcD.

These 3 best experiments lead to 89% sequence coverage for the NISTmAb Lc and 47% for the Hc. For the Lc, 89% is very close to the maximum of 92% achieved with 6 combined experiments (and more). However, the situation is not as optimal for the Hc since the maximum is 70% for 12 and more combined experiments. As depicted in Figure 6, a plateau is rapidly reached for the Lc, although additional experiments always bring new and unique fragments for the Hc. We then sought to make the same comparison but using only HCD, as the use of an unmodified benchtop Orbitrap Exploris would allow. The results are summarized in Figure S13 for both mAbs. The best combination of 3 LC-MS/MS runs with HCD activation corresponds to 2 multiplex and 1 experiment of the highest charge state and leads only to 42% for the Lc and 24% for the Hc. These results clearly show the added value of EThcD for improved sequence coverage.

## 4. CONCLUSION

In this work, we extensively studied for the first time on an LC time scale, the combination of parameters leading to the best sequence coverage for the top-down analysis of light and heavy chains formed from the reduction of intact antibodies. Theses parameters are the precursor charge state (individual and multiplex), the fragmentation type (HCD, CID, EThcD) and the fragmentation level (NCE for HCD, CE for CID and reaction time and NCE for EThcD). In total, 171 LC-MS/MS runs were acquired (including triplicates) for each mAb: NIST and Sigma. Similar behaviors were obtained for both antibodies. Our results indicate that EThcD is mandatory to achieve the highest coverage of both Lc and Hc. HCD alone leads only to half cleavages, which clearly shows that Tribrid Orbitrap mass spectrometers remain to date the best platform for top-down proteomics. We also demonstrate that multiplexing leads to superior results than selecting individual precursor charge states. Multiplexing allows higher sequence coverage to be achieved in a lower number of experiments, which can be of interest in case of limited amount of sample. However, it is worth noting that, even with the best combination of parameters, a complete sequence coverage (100%) is never achieved, which leaves room for improvement. Moreover, the current trend toward an increased diversity of scaffolds in research pipelines of pharmaceutical companies will make this point even more critical in upcoming years.

## AUTHOR INFORMATION

### Author Contributions

The manuscript was written through contributions of all authors. All authors have given approval to the final version of the manuscript.

## ACKNOWLEDGEMENTS

This work was supported by the European Union’s Horizon 2020 Research and Innovation Program under the grant numbers 829157 (TopSpec) and 823839 (EPIC-XS). J.D acknowledges the Association Nationale de la Recherche et de la Technologie for funding his PhD under agreement CIFRE2020/0483. J.D. and J.C.R aknowledge the Region Ile de France (DIM 1HEALTH) and the LABEX IBEID (grant n°ANR-10-LABX-62-IBEID from the Programme d’Investissements d’Avenir) for funding the Eclipse Orbitrap.

## DATA AVAILABILITY STATEMENT

Data are available via ProteomeXchange with identifier PXD038619.

**ORCID**

## Notes

### Competing Interest Statement

The authors have declared no competing interest.

