## Supplementary Information for "A multi-parameter optimization in middle-down analysis of monoclonal antibodies by LC-MS/MS"

**Table S1.** Targeted m/z and corresponding charge states for light (Lc) and heavy (Hc) chains of both mAbs

| mAb | Subunit | Targeted m/z | Charge state |
|---|---|---|---|
| SiLuLite (Sigma mAb) | Lc | 883.35 | 26+ |
| | | 998.45 | 23+ |
| | | 1148.07 | 20+ |
| | | 1350.50 | 17+ |
| | | 883.35; 998.45; 1148.07; 1350.50 | Multiplex |
| | Hc | 885.27 | 57+ |
| | | 970.30 | 52+ |
| | | 1073.41 | 47+ |
| | | 1230.34 | 41+ |
| | | 885.27; 970.30; 1073.41; 1230.34 | Multiplex |
| HzIgG1 (NISTmAb) | Lc | 857.53 | 27+ |
| | | 964.61 | 24+ |
| | | 1102.28 | 21+ |
| | | 1361.41 | 17+ |
| | | 857.53; 964.61; 1102.28; 1361.41 | Multiplex |
| | Hc | 912.97 | 56+ |
| | | 1002.37 | 51+ |
| | | 1111.22 | 46+ |
| | | 1277.73 | 40+ |
| | | 912.97; 1002.37; 1111.22; 1277.73 | Multiplex |

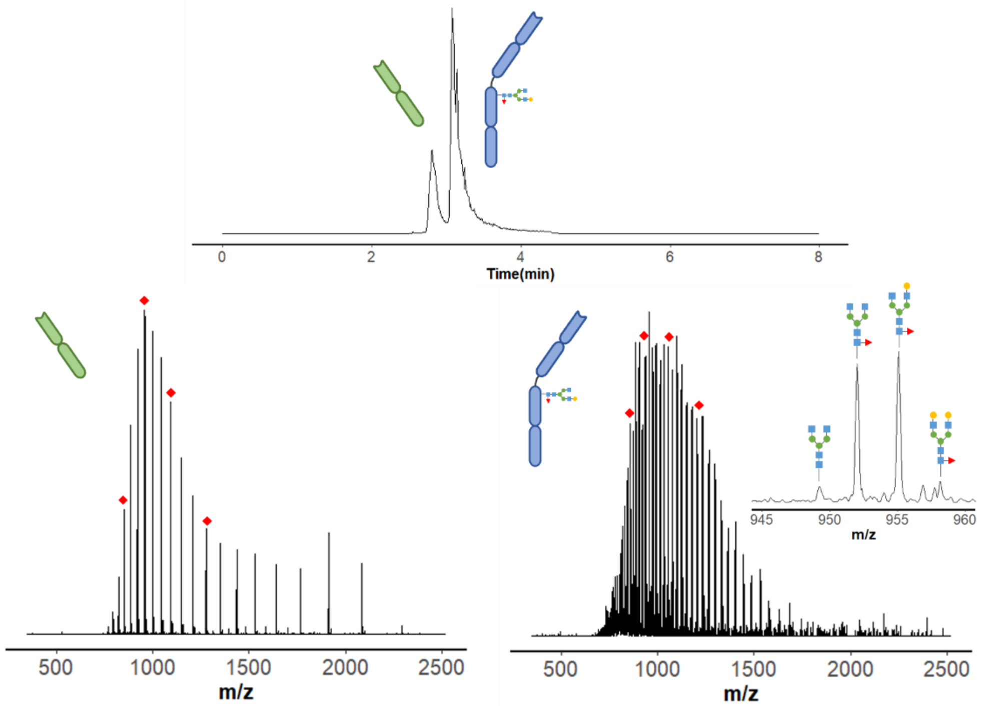


**Figure S1.** Chromatogram of Sigma mAb Lc and Hc (top), MS spectra of Lc (bottom left) and Hc (bottom right). Precursor charge states selected for MS/MS analysis are depicted as red diamonds in MS spectra.

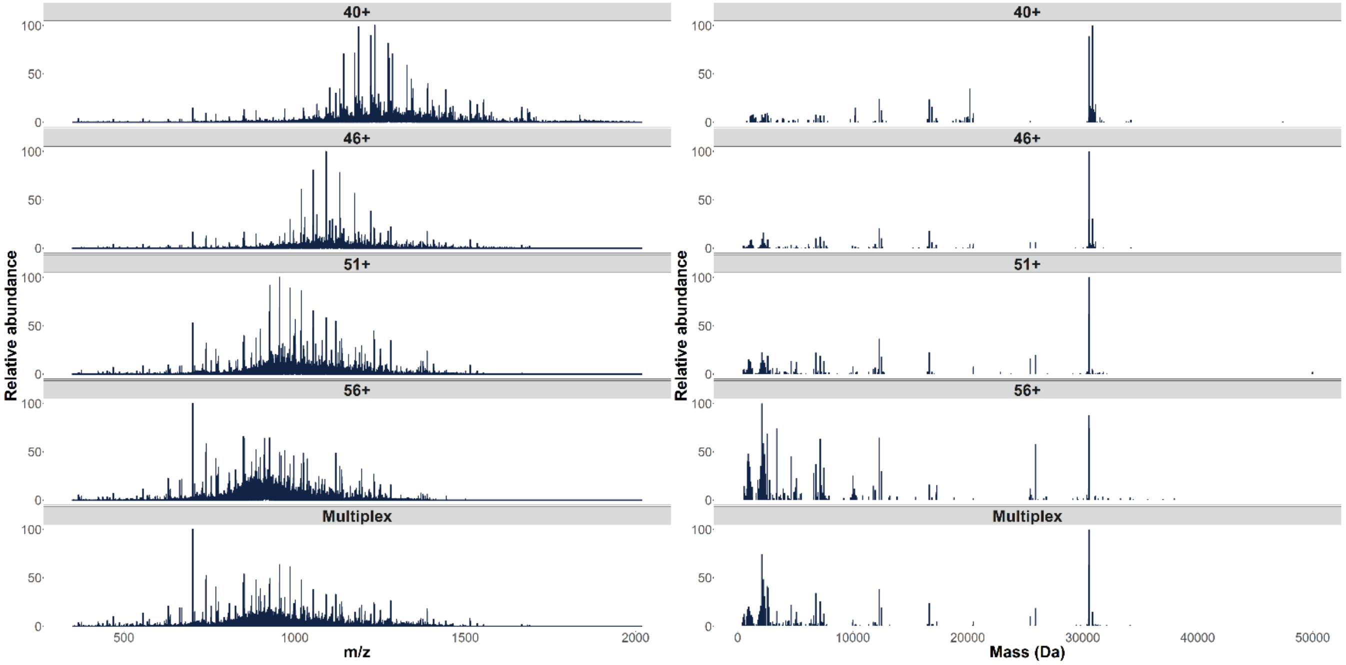


**Figure S2.** HCD MS/MS spectra with NCE25% for each selected charge state of NISTmAb Hc before (left) and after deconvolution (right). From top to bottom: 40+, 46+, 51+, 56+ and multiplex

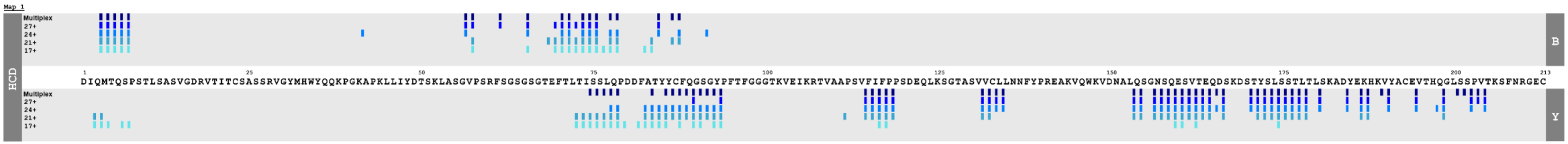


| Precursor charge state | Multiplex | 27+ | 24+ | 21+ | 17+ |
|---|---|---|---|---|---|
| Residue cleavages (%) | 34 | 28 | 32 | 29 | 17 |
| Number of matching fragments | 78 | 59 | 73 | 71 | 48 |

**Figure S3.** TDFragMapper fragmentation map of NISTmAb Lc obtained when varying the precursor charge state in HCD with NCE25%

**Table S2.** Comparison of residue cleavages between multiplex and the sum of individual precursors for the NISTmAb Hc and percentages of cleavages from individual fragmentations contained in multiplex experiments

| **Fragmentation experiment** | **Cumulated residue cleavages (%)** | | **Cleavages from individual precursors observed in multiplex (%)** |
|---|---|---|---|
| | Multiplex | Individual precursors | |
| **HCD 15%** | 11 | 21 | 30 |
| **HCD 20%** | 13 | 20 | 48 |
| **HCD 25%** | 13 | 19 | 55 |
| **ET2hcD5** | 21 | 34 | 35 |
| **ET2hcD15** | 21 | 31 | 36 |
| **ET5hcD5** | 24 | 26 | 77 |
| **ET5hcD15** | 27 | 24 | 82 |
| **ET10hcD10** | 22 | 18 | 84 |
| **ET10hcD15** | 24 | 16 | 90 |

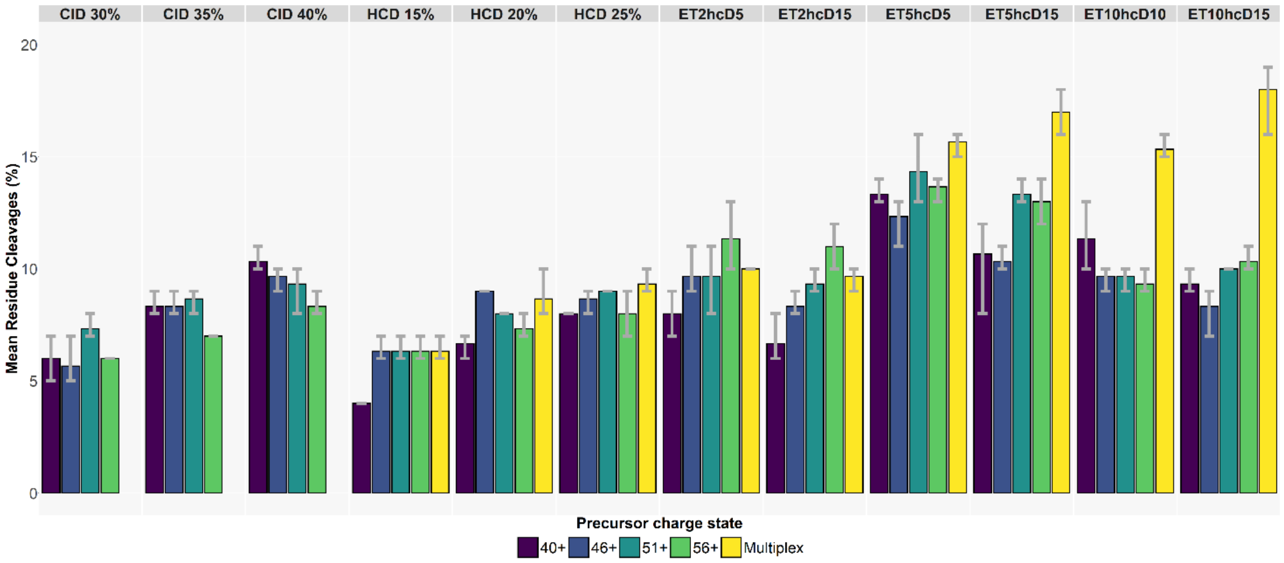


**Figure S4.** Mean residue cleavages (triplicates) for each selected precursor charge states of the NISTmAb Hc across the different fragmentation experiments

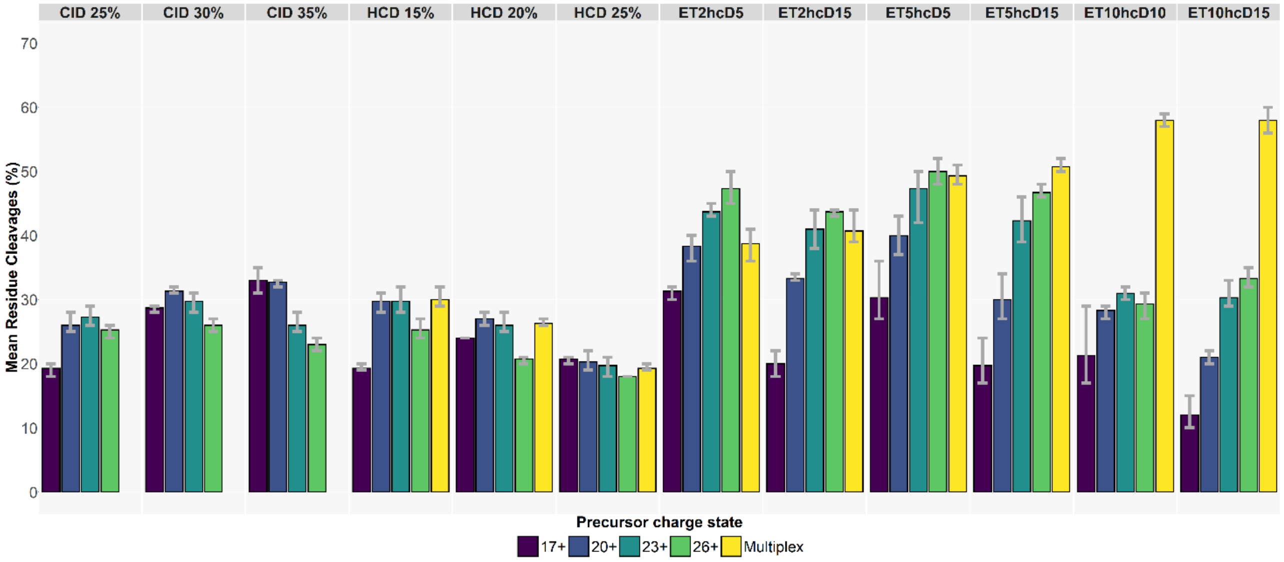


**Figure S5.** Mean residue cleavages (triplicates) for each selected precursor charge states of the Sigma mAb Lc across the different fragmentation experiments

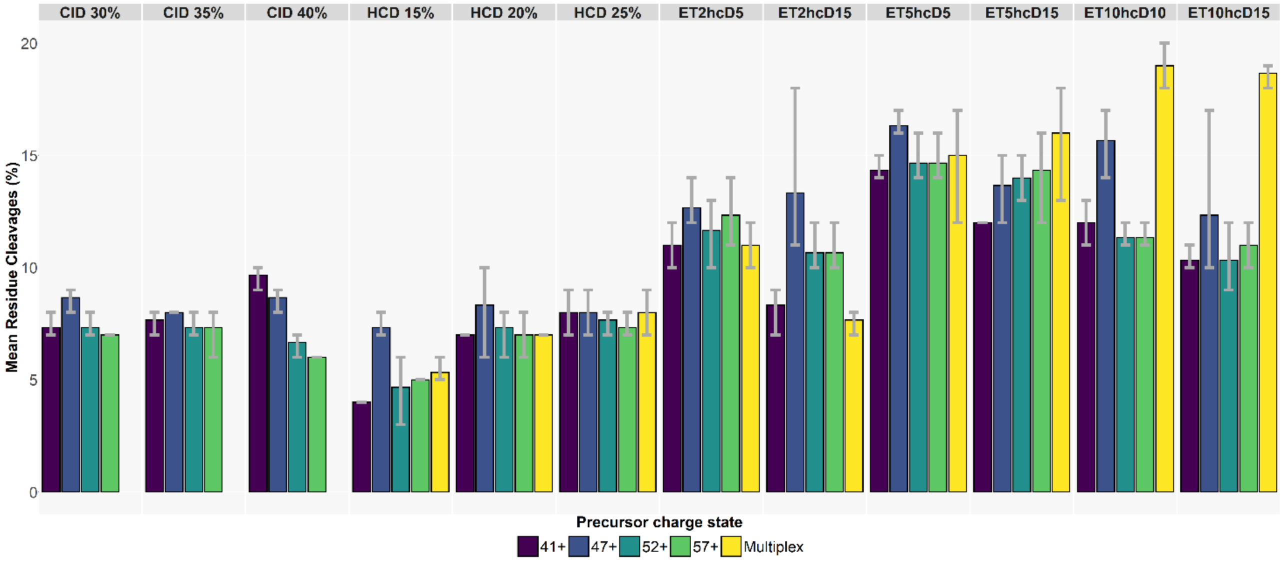


**Figure S6.** Mean residue cleavages (triplicates) for each selected precursor charge states of the Sigma mAb Hc across the different fragmentation experiments

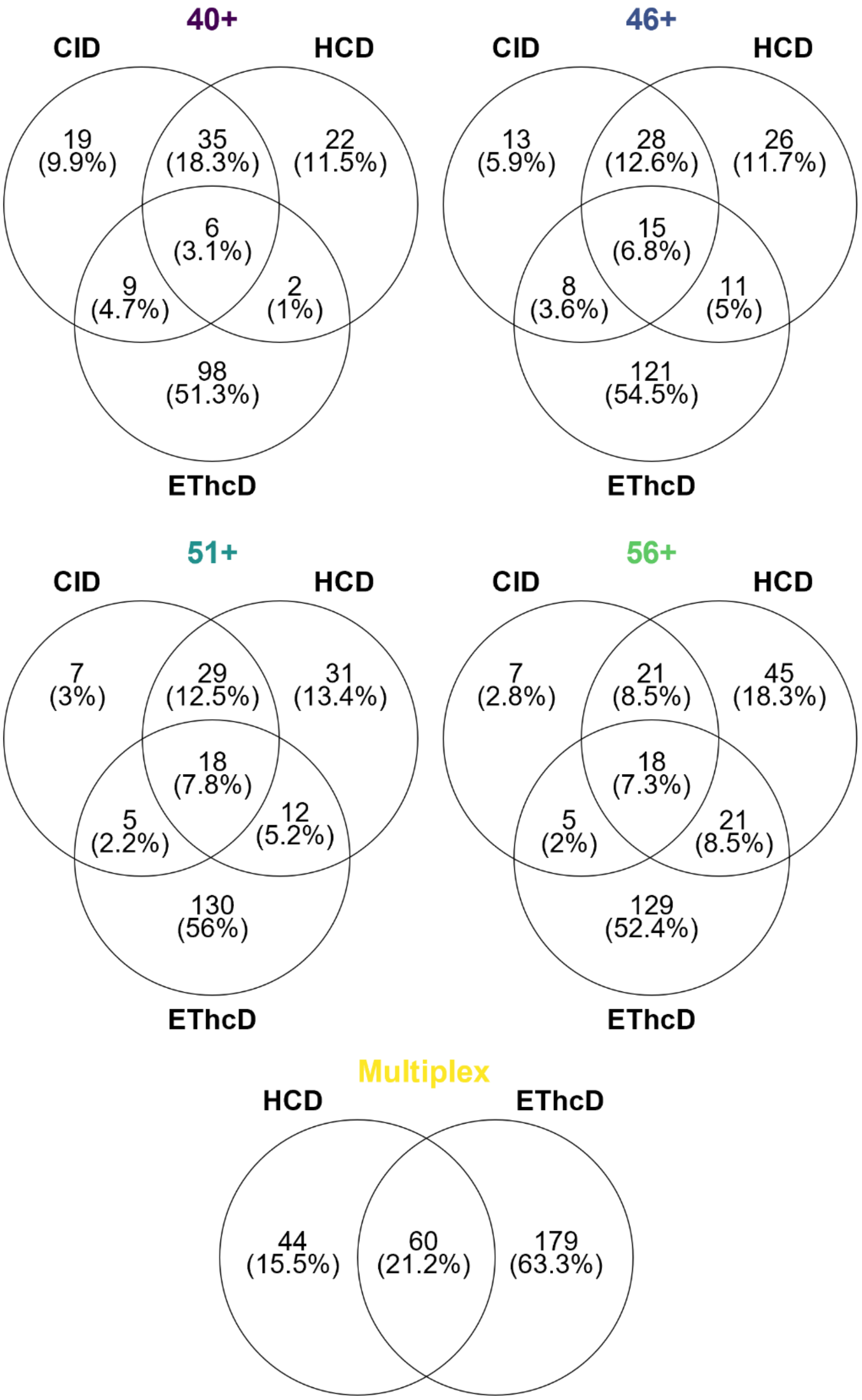


**Figure S7.** Venn diagrams of cleavages generated in CID, HCD and EThcD for every targeted precursor charge state and multiplex for the NISTmAb Hc

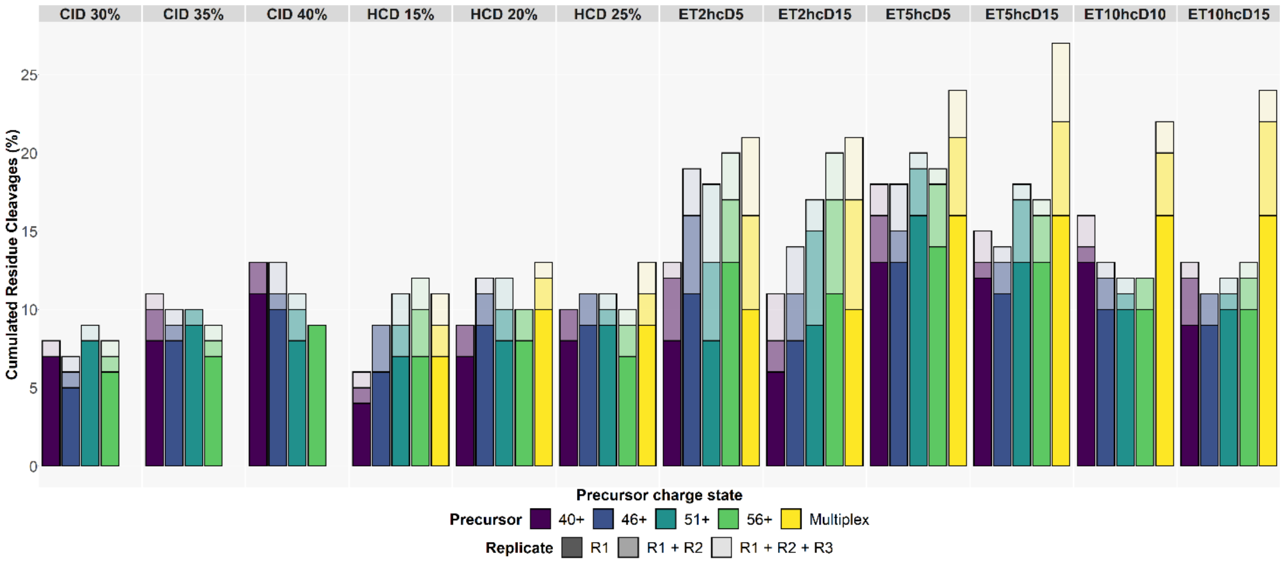


**Figure S8.** Summed residue cleavages over 3 replicates for each selected precursor charge states of the NISTmAb Hc across the different fragmentation experiments

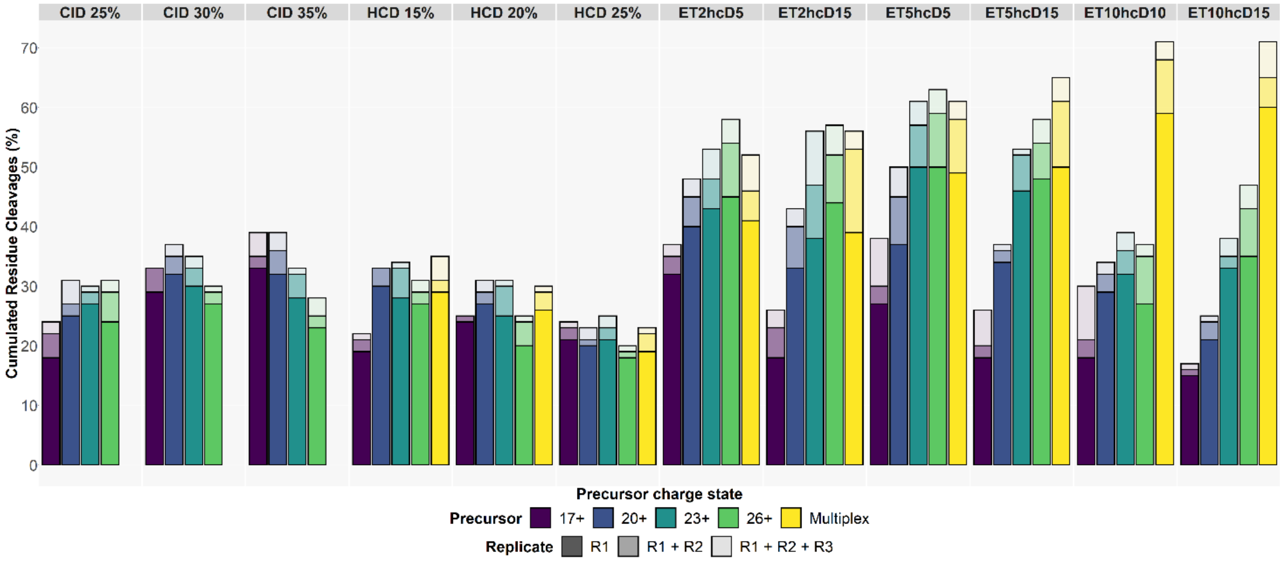


**Figure S9.** Summed residue cleavages over 3 replicates for each selected precursor charge states of the Sigma mAb Lc across the different fragmentation experiments

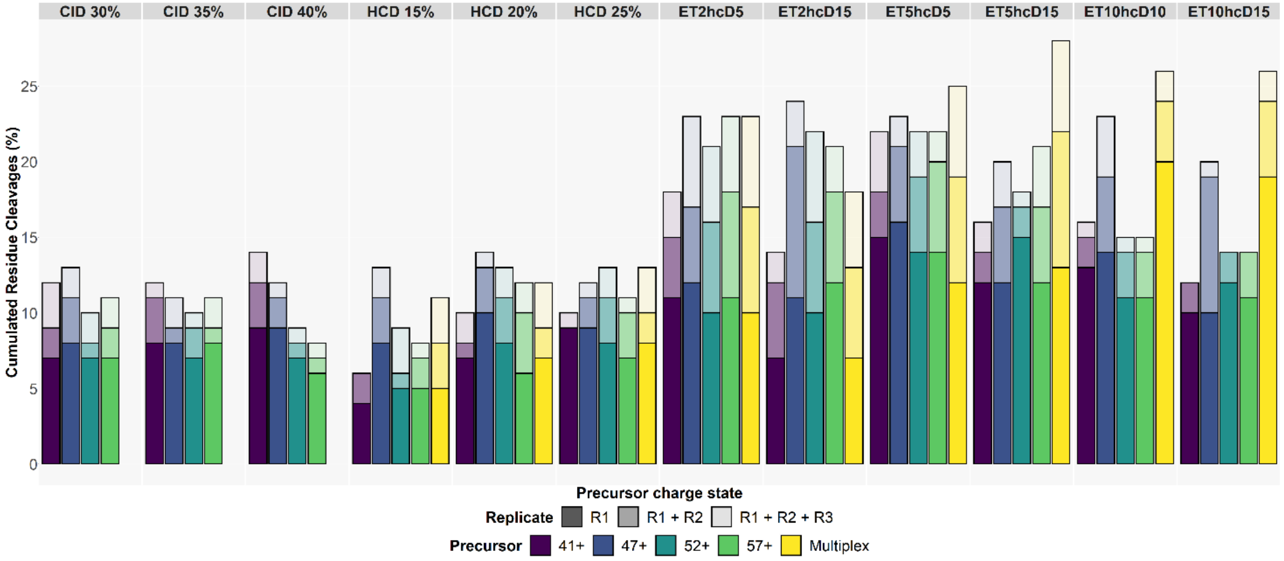


**Figure S10.** Summed residue cleavages over 3 replicates for each selected precursor charge states of the Sigma mAb Hc across the different fragmentation experiments

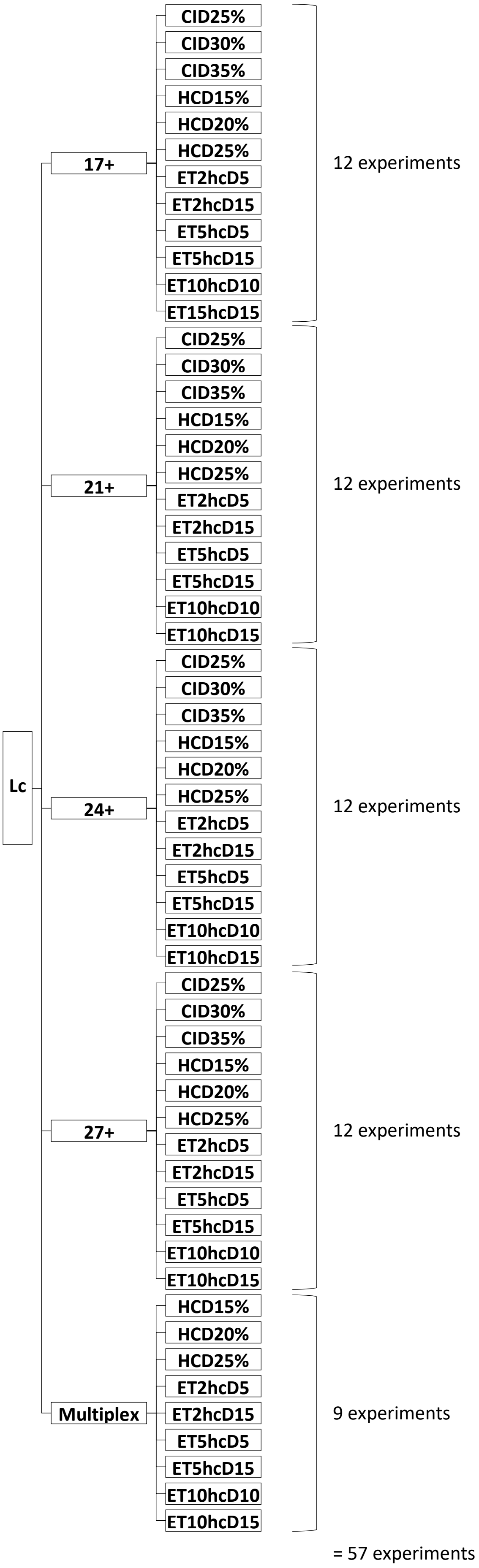


$$Total\ number\ of\ combinations = \sum_{i=1}^{57} \binom{57}{i} = 2^{57}$$

**Figure S11.** LC-MS/MS experiments tree for the NISTmAb Lc

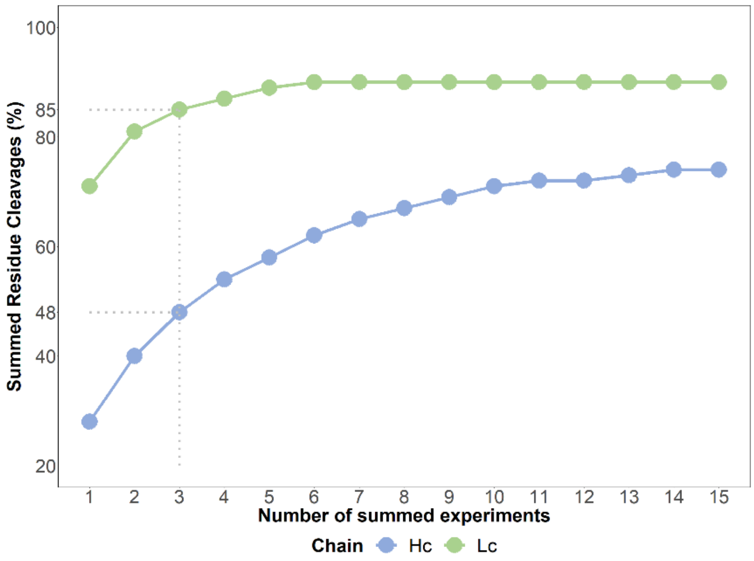


**Figure S12.** Maximum residue cleavage (%) obtained when combining diverse MS/MS experiments for the Sigma mAb Lc and Hc

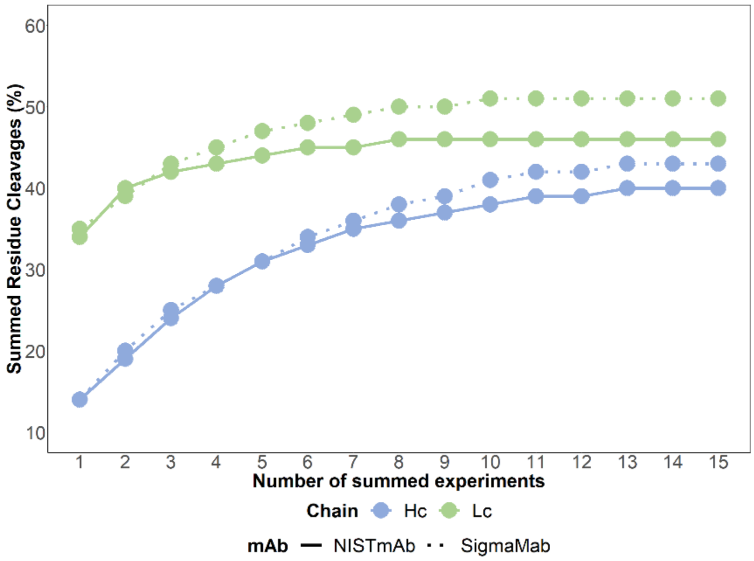


**Figure S13.** Maximum residue cleavage (%) obtained when combining only HCD MS/MS experiments for Lc and Hc of both mAbs
